# Knockout of E-cadherin in adult mouse epithelium results in emphysema and airway disease

**DOI:** 10.1101/2021.07.18.452342

**Authors:** Baishakhi Ghosh, Jeffrey Loube, Shreeti Thapa, Erin Capodanno, Saborny Mahmud, Mirit Girgis, Si Chen, Kristine Nishida, Linyan Ying, Carter Swaby, Ara Wally, Debarshi Bhowmik, Michael Zaykaner, Wayne Mitzner, Venkataramana K. Sidhaye

**Author notes:** Corresponding author Dr. Venkataramana Sidhaye, Contact No. Department of Pulmonary and Critical Care Medicine; Shanghai East Hospital; Tongji University; Shanghai, 200120, China (Si Chen). Department of Respiration, Children’s Hospital of Chongqing Medical University, Chongqing, China (Linyan Ying). **Author contributions** AW, BG, CS, DB, EC, KN, LY, MG, MZ, SC, SM, and ST executed all experiments. BG contributed to the experimental design, implementation, prepared figures, data analysis, data interpretation and drafting the manuscript. JL performed lung function test on mice under the supervision of WM. VKS is the principal investigator, who conceived the idea, led the study design, data interpretation, and edited the manuscript. All authors have read and approved the manuscript.

## Abstract

Chronic obstructive pulmonary disease (COPD) is a devastating lung disease, characterized by a progressive decline in lung function, alveolar loss (emphysema), and airflow limitation due to excessive mucus secretion (chronic bronchitis), that can occur even after the injurious agent is removed. It is slated to rise to the 3^rd^ leading cause of death due to chronic disease by 2030 globally, and the 4^th^ leading cause of death due to chronic disease in the USA. While there is substantial evidence indicating loss of E-cadherin in the lung epithelium of patients with COPD, it is not known if this is causal to the disease. We investigated if loss of E-cadherin can result in lung disease using in both in vitro models of primary, differentiated human cells and in mouse models. Using a cell type-specific promoter using Cre/LoxP mice system to knock-out E-cadherin in ciliated and alveolar epithelial cell (Type 1 and Type 2) populations in adult mouse models, we determined that loss of E-cadherin caused airspace enlargement, as well as increased airway hyperresponsiveness indicating that it does have a causative role in causing COPD. Strategies to upregulate *CDH1* (encodes for E-cadherin) in CHBEs and cigarette-smoke injured NHBEs can rescue the dysfunctional epithelium.

## Introduction

Chronic obstructive pulmonary disease (COPD) is a devastating lung disease that is the slated to rise to the 3^rd^ leading cause of death due to chronic disease by the 2030 globally, and 4^th^ leading cause of death due to chronic disease in USA (1–3). It characterized by progressive decline in lung function, persistent alveolar loss (emphysema), and airflow limitation due to excessive mucus secretion (chronic bronchitis), that can occur even after the injurious agent is removed. The etiologic association with noxious insults such as cigarette smoke (CS) and COPD is clear, with CS quantitatively altering both structure and function of the lung epithelium (4–8). The lung epithelium is the site of initial contact with inhaled agents and establishes a protective physical barrier with robust cell-cell contacts to limit access between the external environment and sub-epithelial tissues. The epithelial barrier function is maintained by formation of adherens junctions (AJs), tight junctions (TJs) and desmosomes (9). This barrier is compromised in COPD and CS injured epithelium due to decreased barrier function with increased permeability and preserved apical-basolateral polarity, reduced percentage of moving cilia with decreased ciliary beat frequency (CBF) and an unjammed epithelium (5, 10–12). E-cadherin is an adherens junctional protein which mediates architecture of the epithelia, paracellular permeability, and suppresses intracellular signaling pathway, regulates epithelial activation, proliferation, and differentiation (13, 14).

We and others have previously reported that chronic exposure to CS to primarily human bronchial epithelial cells in an air-liquid interface reduces E-cadherin and patients with COPD have less E-cadherin (4–6, 15–17). In addition, decreases in E-cadherin are associated with the development of COPD (18–22) with COPD tissues displaying loss of E-cadherin associated with de-differentiation of epithelium with evidence of subepithelial fibrosis (20). Moreover, there is evidence that lack of E-cadherin in histology of human lung specimens of COPD suggests decreased or degraded E-cadherin in emphysematous regions, indicating its role to maintain the integrity of pulmonary epithelium (21, 22). While there is evidence that loss of lung epithelial E-cadherin affects lung development (23), it is not known if loss of E-cadherin is sufficient to cause the tissue damage seen in COPD. As such, conclusive evidence for a causal role of loss in E-cadherin in adult lungs in tissue modeling is lacking.

In this study, we investigated if loss of E-cadherin is causal to disease using primary cell in vitro models and in mouse models. We assessed if loss of E-cadherin can cause epithelial dysfunction in primary non-diseased human bronchial epithelium (NHBEs) similar to that seen in human bronchial epithelial cells derived from patients with COPD (CHBEs). In addition, we implemented a cell type-specific promoter using Cre/LoxP mice system to knock-out E-cadherin in ciliated and alveolar epithelial cell (Type 1 and Type 2) populations in adult mouse models to further investigate its causative role in lung injury by quantifying lung function and lung morphometry. Moreover, we investigated if strategies to upregulate *CDH1* (encodes for E-cadherin) in CHBEs and CS injured NHBEs can rescue epithelium dysfunction.

## Results

### E-cadherin knock down in non-diseased airway epithelia causes epithelial dysfunction

To assess if knockdown of E-cadherin within the non-diseased airway epithelium affects lung injury, adenovirus-based shRNA targeting the *CDH1* (Ad-GFP-U6-shRNA, shCDH1) and Adenovirus-based scrambled shRNA with GFP (Ad-GFP-U6-shRNA, GFP) were used in NHBEs at week 4 to 6 ALI to evaluate epithelial dysfunction. We observed ∼50% of E-cadherin knockdown in NHBEs as observed under fluorescence at 20X magnification, and western blot assay (Fig. 1A – C). This knockdown significantly decreased barrier function (Fig. 1D) with increasing trend of monolayer permeability (Fig. 1E), contributing to an unjammed epithelium (Fig. 1F). Interestingly, the E-cadherin knockdown had no effect on % cilia moving and CBF (Fig. 1G). Also, NHBE with E-cadherin knockdown showed lower expression of *CDH1* (encodes for E-cadherin) and higher expression of EMT markers such as *CDH2, SNAI2, SLUG*, and TWIST2 (Fig. 1H – L).

**Figure 1.**
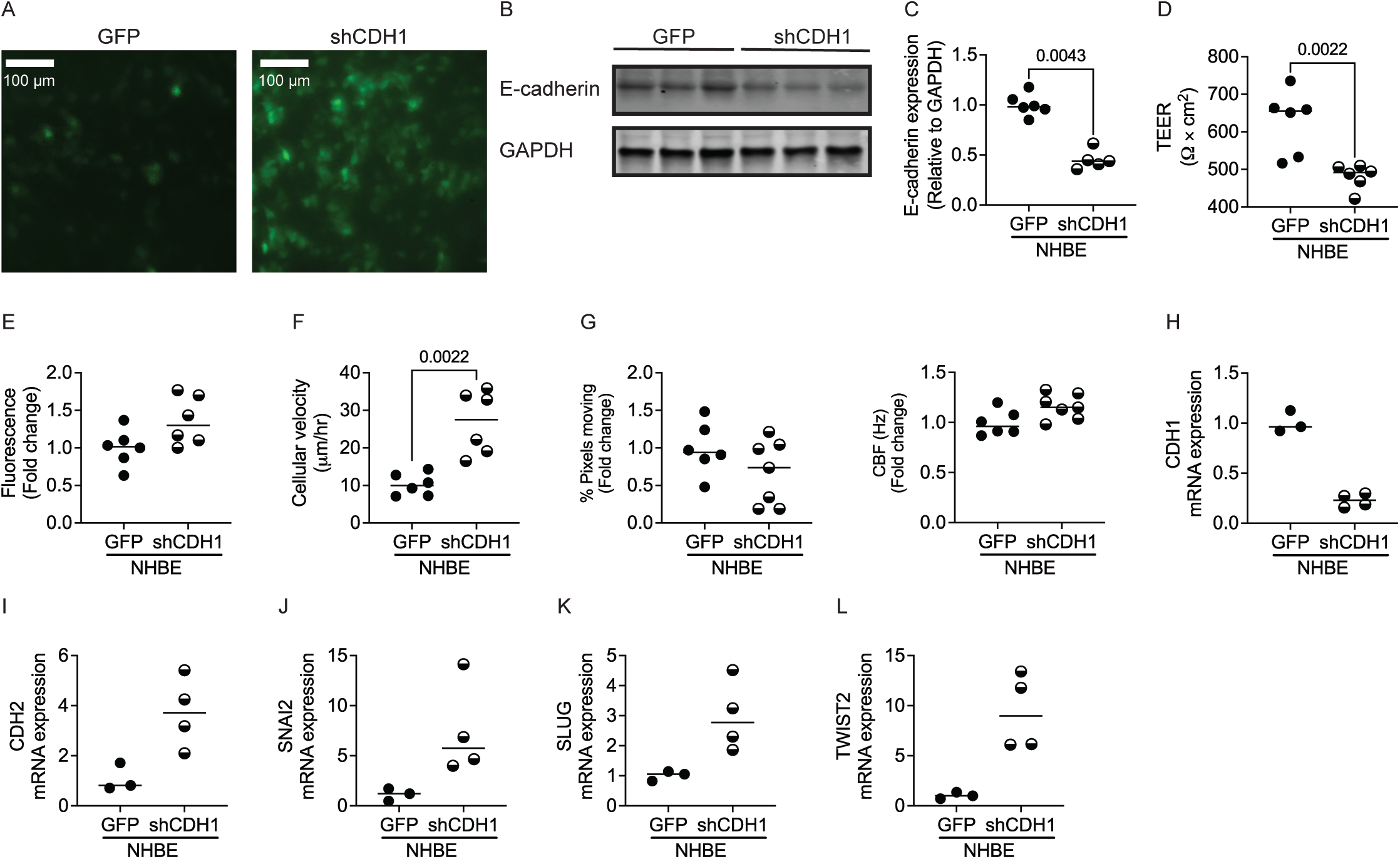
Knockdown of E-cadherin in non-diseased airway epithelia (NHBE) causes epithelial dysfunction. (A) GFP 20X view (scale bar 100 µm), (B) Represent blot of E-cadherin and GAPDH, and (C) quantified blot of E-cadherin relative to GAPDH, after 48h transfection with Control Adenovirus-based scrambled shRNA with GFP (Ad-GFP-U6-shRNA, GFP), and Adenovirus to knock out E-cadherin (Ad-GFP-U6-h-CFL1-shRNA, shCDH1). Assessment of epithelial functional phenotypes such as (D) barrier function, (E) permeability, (F) cellular velocity, and (G) percentage moving cilia (left panel) and ciliary beat frequency (right panel), of NHBE transfected with GFP/shCDH1. Assessment of (H) CDH1 as epithelial marker, and (I) CDH2, (J) SNAI2, (K) SLUG, and (L) TWIST2 as mesenchymal markers of NHBE transfected with GFP/shCDH1. Data is representative of 2 donors and 2 to 3 inserts per donor. *P* < 0.05 were considered statistically significant.

### Loss of E-cadherin in mice tracheal epithelial cells (mTECs) causes airway epithelial dysfunction

To evaluate if loss of E-cadherin can contribute to airway epithelial dysfunction phenotypes, we have isolated the mTEC from *Cdh1*^*fl/fl*^ mice *ex vivo* at 14 days ALI and treated with adenoviruses Ad5CMVeGFP (Ctrl) or Ad5CMVCre-eGFP (Cre). We observed that loss of E-cadherin contributed to decreased barrier function (Fig. 2A) and an unjammed epithelium as observed with increased cellular velocity (Fig. 2B), without any effect on % cilia moving and CBF (Fig. 2 C). We also quantified the *CDH1* (encodes for E-cadherin) mRNA and protein expression of E-cadherin in *ex vivo*. There was ∼50% reduced *Cdh1* mRNA expression (Fig. 2D) and ∼75% E-cadherin protein downregulation (Fig. 2E & F).

**Figure 2.**
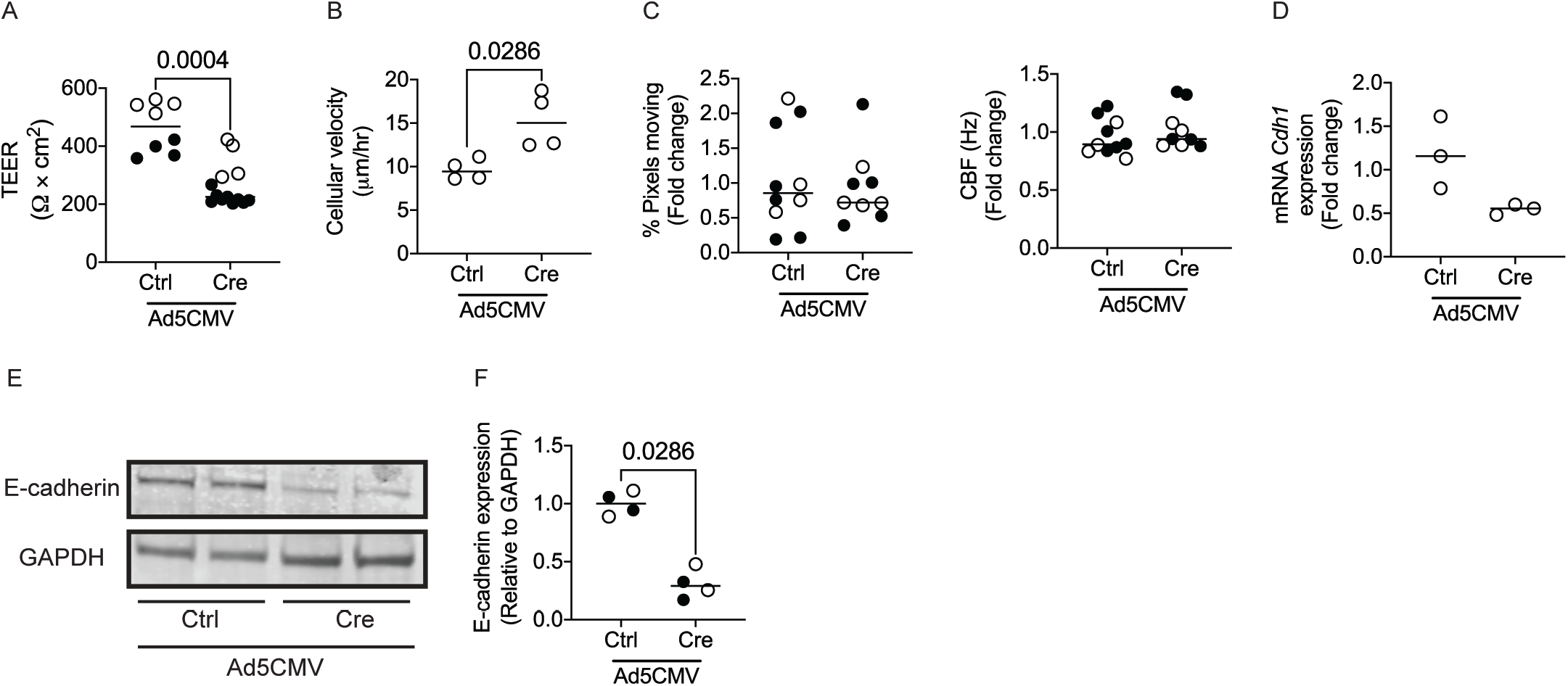
Loss of E-cadherin in mice tracheal epithelial cells (mTECs) causes airway epithelial dysfunction. Assessment of epithelial functional phenotypes such as (A) barrier function, (B) cellular velocity, (C) percentage moving cilia (left panel), and ciliary beat frequency (right panel) of mTECs transfected with either Adenovirus Ad-5CMVeGFP (Cntrl) and Ad5CMVCre-eGFP (Cre). Loss of E-cadherin was assessed by (D) mRNA expression of *Cdh1* (encodes for E-cadherin), (E) representative western blot of E-cadherin and GAPDH and (F) quantification of E-cadherin expression relative to GAPDH. Data is representative from cells derived from 12 mice, 3 to 12 inserts. *P* < 0.05 were considered statistically significant.

### In vivo E-cadherin knockdown in mice lung contributes to increased lung morphometry and decreased lung function

To determine if E-cadherin has a causal role in airway dysfunction and parenchymal modeling, we have performed bimonthly intratracheal instillations of adenoviruses Ctrl or Cre on *Cdh1*^*fl/fl*^ mice and lung morphometry and lung function was assessed. Exposure to adeno-Cre causes reduction in E-cadherin in mouse lungs as quantified by immunofluorescence after 10 days (Fig. 3A), and this was consistent throughout the instillations period (Fig. 3B & C). After 1-month instillation, the Hemotoxylin and eosin (H&E) staining showed increased terminal airway/airspace enlargement (Fig. 3D) with higher mean linear intercept (MLI, Lm) in Cre delivery group (Fig. 3E), indicating loss of E-cadherin in lung epithelium induces emphysematous injuries. Also, the adeno-Cre delivered group had significantly higher total lung capacity (Fig. 3F) and vital capacity (Fig. 3G), without any difference in the residual volume (Fig. 3H). The lung compliance, which measures the ability of a lungs to stretch and expand showed an ascending trend, but not statistically significant (Fig. 3I).

**Figure 3.**
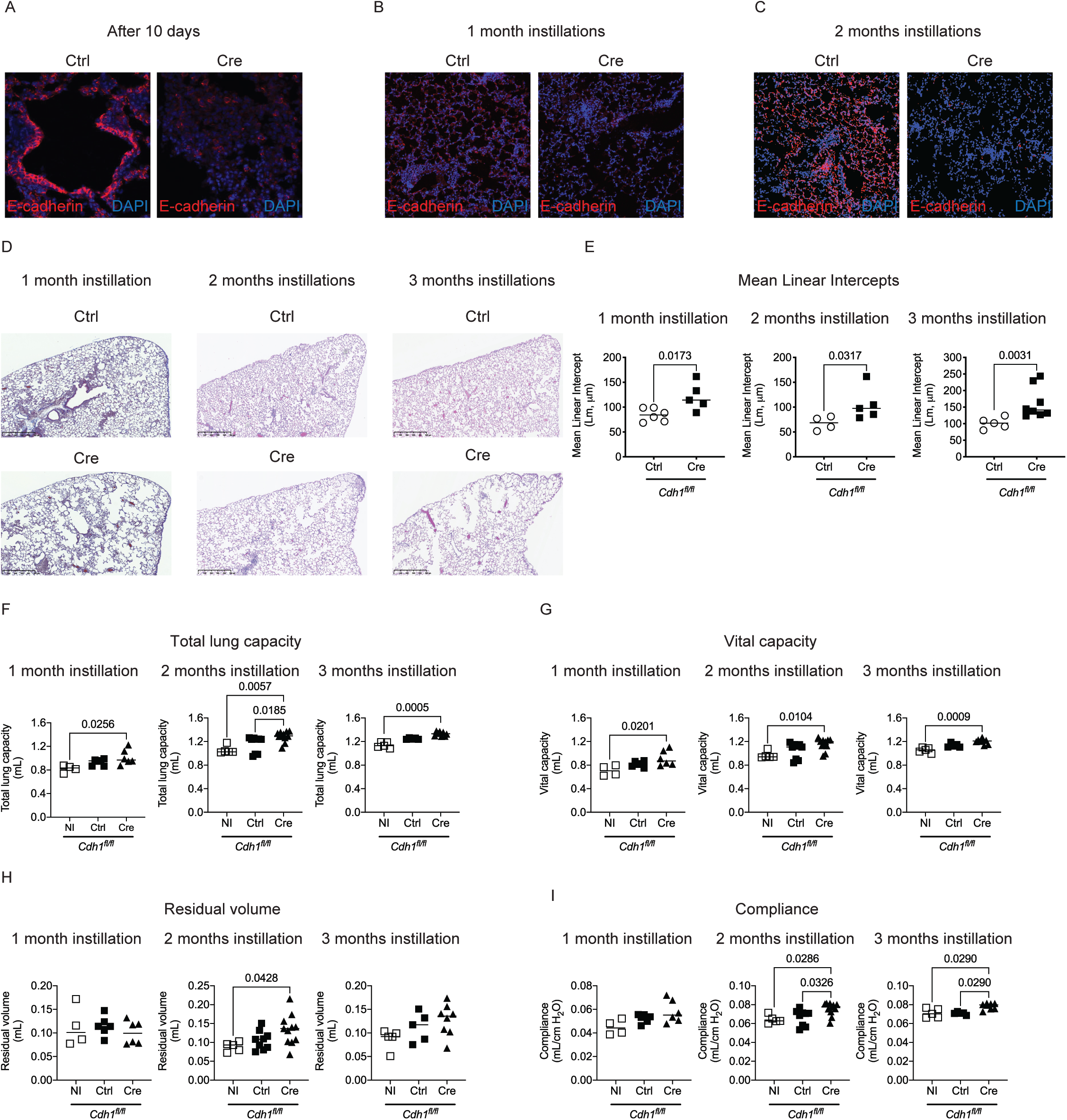
In vivo E-cadherin knockdown in mice lung contributes to increased lung morphometry and decreased lung function. (A) Immunofluorescence of E-cadherin (Red) and DAPI (Blue) on mice lung parenchyma after 10 days of tracheal instillations of adeno-Ctrl (Ad-5CMVeGFP) or Cre (Ad5CMVCre-eGFP) instillations. Immunofluorescence of E-cadherin (Red) and DAPI (Blue) on mice lung parenchyma at 10X after (B) 1 month and (C) 2 months of tracheal instillations of adeno-Ctrl or Cre. (D) Representative H & E staining of mice lung parenchyma and (E) quantified mean linear intercepts (Lm) after 1 to 3 months with adeno-Ctrl or Cre after 1 month (left panel) or 2 months (middle panel) or 3 months (right panel). Assessment of (F) total lung capacity, (G) vital capacity, (H) residual volume, and (I) compliance after 1 to 3 months instillations with adeno-Ctrl or Cre. Data is representative from 4 to 11 mice. *P* < 0.05 were considered statistically significant.

To better characterize the observed emphysema and fibrosis observed in 1 month instillation, we further increased the timepoints for the instillations. The H & E staining showed patchy regions and increased terminal airway/airspace enlargement with increased MLI as observed in Cre delivery group as observed in 1-month instilled mice group (Fig. 3D & Fig. 3E). We observed significantly higher total lung capacity and vital capacity with higher residual volume and lung compliance in 2-month and 3-month instillations (Fig. 3F-I). These data suggest evidence of injury with emphysema and fibrosis in the mice lung.

### E-cadherin knock down in specific cell type of lung epithelium

Since E-cadherin regulates structure and function of the lung epithelium, we further assessed if loss of E-cadherin in a specific cell types of particularly ciliated cells, and alveolar type I (AT 1) and II (AT2) cells can contribute to the injuries as observed in whole lung model. To answer this, we have generated tamoxifen inducible specific mice to knock out E-cadherin in ciliated cells *Cdh1*^*fl/fl*^(*Foxj1-CreERT*^*2*^), AT1 cells *Cdh1*^*fl/fl*^(*Ager-CreER*^*T2*^), and AT2 cells *Cdh1*^*fl/fl*^(*Sftpc-CreERT*^*2*^).

To knockdown E-cadherin in ciliated cells, *Cdh1*^*fl/fl*^_*Hom/Het*_*Foxj1*_*Het*_ received Tamoxifen chow (TAM) diet, which were compared with *Cdh1*^*fl/fl*^_*Hom/Het*_*Foxj1*_*Het*_ receiving normal chow (ND) diet and *Cdh1*^*fl/fl*^_*Hom/Het*_*Foxj1*_*WT*_ receiving TAM diet as control group. The lung function parameters (total lung capacity, vital capacity, residual volume, and compliance) and MLI were not affected in *Cdh1*^*fl/fl*^_*Hom/Het*_*Foxj1*_*Het*_ receiving TAM diet compared with *Cdh1*^*fl/fl*^_*Hom/Het*_*Foxj1*_*Het*_ receiving ND and *Cdh1*^*fl/fl*^_*Hom/Het*_*Foxj1*_*WT*_ receiving TAM (Fig. 4A – D & F). Interestingly, we observed that *Cdh1*^*fl/fl*^_*Hom/Het*_*Foxj1*_*Het*_ receiving TAM diet had significantly increased airway hyperresponsiveness (AHR) than *Cdh1*^*fl/fl*^_*Hom/Het*_*Foxj1*_*Het*_ receiving ND and *Cdh1*^*fl/fl*^_*Hom/Het*_*Foxj1*_*WT*_ receiving TAM (Fig. 4E), indicating that E-cadherin knockdown in ciliated cells contributes in an increased AHR.

**Figure 4.**
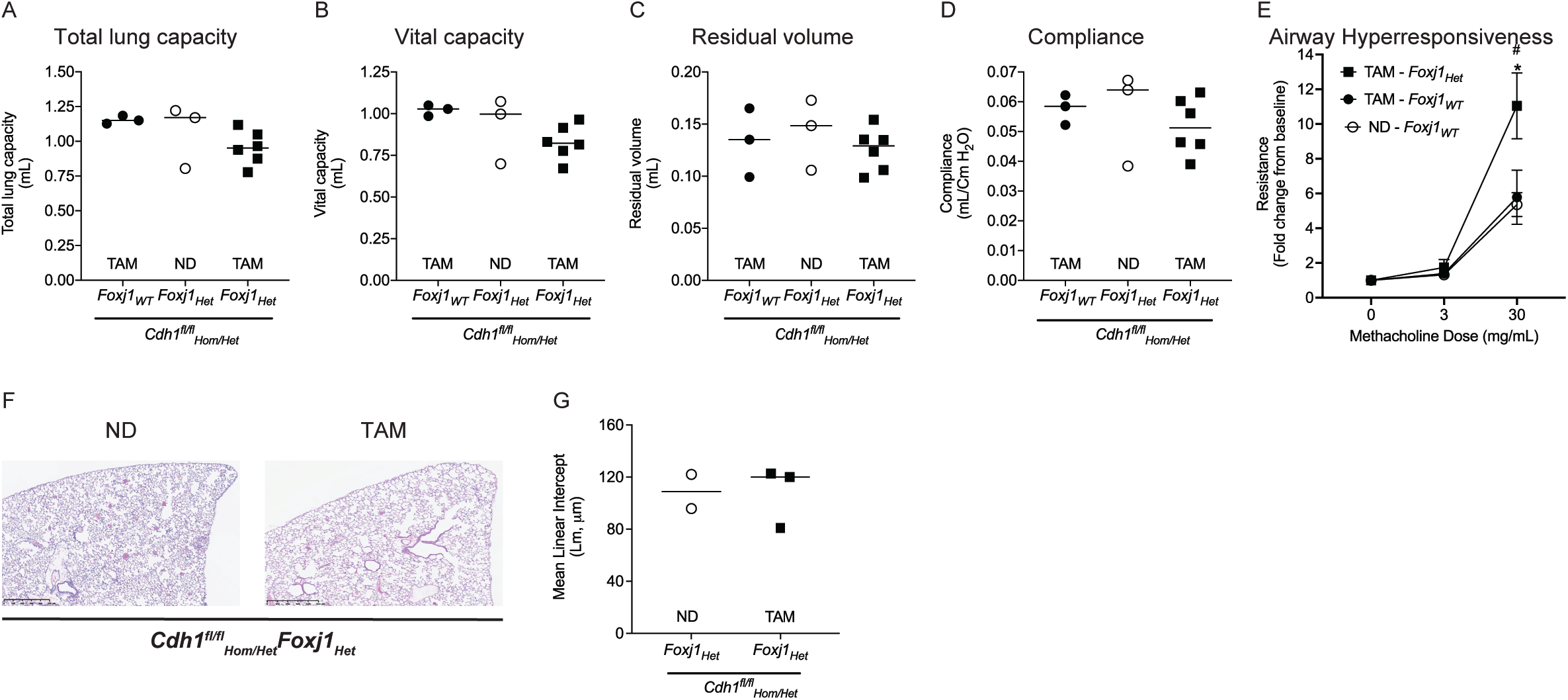
*In vivo* E-cadherin knockdown in the ciliated cells of mice lung contributes to increased airway hyperresponsiveness. To knockdown E-cadherin in the ciliated cells of mice lung, *Cdh1*^*fl/fl*^_*Hom/Het*_*Foxj1*_*Het*_ mice were fed tamoxifen (TAM) for 30 days. These were compared to *Cdh1*^*fl/fl*^_*Hom/Het*_*Foxj1*_*Het*_ mice receiving normal chow diet (ND) and *Cdh1*^*fl/fl*^_*Hom/Het*_*Foxj1*_*WT*_ receiving TAM control mice. Assessment of (A) total lung capacity, (B) vital capacity, (C) residual volume, (D) compliance and (E) airway hyperresponsiveness in ciliated cells of mice lung. (F) Representative H & E staining of mice lung parenchyma and (G) quantified mean linear intercepts (Lm) after E-cadherin knockdown in ciliated cells. Data is representative from 3 to 6 mice. *P* < 0.05 were considered statistically significant.

As the AT1 cells cover 95% of the alveolar cells (24), we further evaluated the consequences of knocking down E-cadherin in these cells. To knockdown E-cadherin in AT1 cells, *Cdh1*^*fl/fl*^_*Hom/Het*_*Ager*_*Het*_ received TAM diet which were compared with *Cdh1*^*fl/fl*^_*Hom/Het*_*Ager*_*Het*_ receiving ND diet and *Cdh1*^*fl/fl*^_*Hom/Het*_*Ager*_*WT*_ receiving TAM as control group. The total lung capacity, vital capacity, and residual volume were significantly reduced in *Cdh1*^*fl/fl*^_*Hom/Het*_*Foxj1*_*Het*_ receiving TAM compared to the control group (Fig. 5A – C). However, the compliance of the lung was not affected due to the knockdown (Fig. 5D). Also, we did not observe any difference in MLI based on lung histology analysis (Fig. 5E & F), indicating E-cadherin knockdown in AT1 cells has a minimal role in a lung injury as observed in whole lung knockdown. Interestingly, we observed an increase in mRNA expression of mesenchymal and collagen markers, without an overall change in E-cadherin expression (Fig. 5G – L). The Masson trichrome stain also show an increase in blue staining around the airways of AT1 cells with knocked down E-cadherin, indicating fibrosis (Fig. 5M).

**Figure 5.**
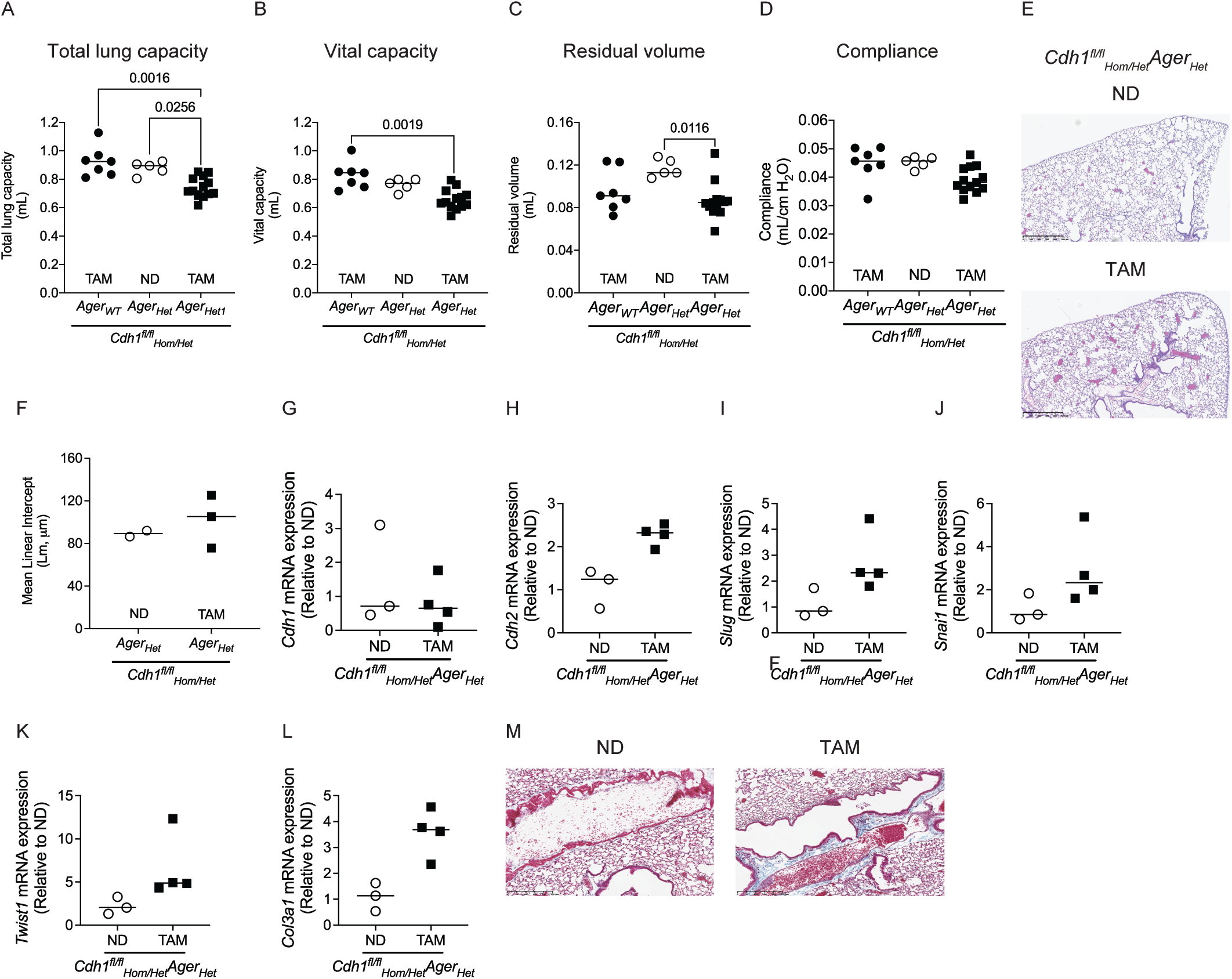
*In vivo* E-cadherin knockdown in the alveolar type I (AT1) cells of mice lung contributes to a decrease in lung capacity. To knockdown E-cadherin in the AT1 cells of mice lung, *Cdh1*^*fl/fl*^_*Hom/Het*_*Ager*_*Het*_ mice were fed tamoxifen (TAM) for 30 days. These were compared to *Cdh1*^*fl/fl*^_*Hom/Het*_*Foxj1*_*Het*_ mice receiving normal chow diet (ND) and *Cdh1*^*fl/fl*^_*Hom/Het*_*Foxj1*_*WT*_ receiving TAM control mice. Assessment of (A) total lung capacity, (B) vital capacity, (C) residual volume, and (D) compliance in E-cadherin knockdown in AT1 cells of mice lung. (E) Representative H & E staining of mice lung parenchyma and (F) quantified mean linear intercepts (Lm) after E-cadherin knockdown in AT1 cells. Comparison of basal mRNA expression of epithelial – (G) Cdh1, mesenchymal – Cdh2 (H), Slug (I), and Snai1 (J) and collagen (L) Col3a1 markers after E-cadherin knockdown in AT1 cells. (M) Representative Masson Trichrome staining of mice lung parenchyma of E-cadherin knockdown AT1 cells. Data is representative from 3 to 6 mice. *P* < 0.05 were considered statistically significant.

To knockdown E-cadherin in AT2 cells, *Cdh1*^*fl/fl*^_*Hom/Het*_*Spc*^*Cre*^ received TAM diet which were compared with *Cdh1*^*fl/fl*^_*Hom/Het*_*Spc*^*Cre*^ receiving ND as control group. The total lung capacity, vital capacity, and compliance were significantly increased in *Cdh1*^*fl/fl*^_*Hom/Het*_*Spc*^*Cre*^ receiving ND (Fig. 6 A, B & D), with no differences in residual volume (Fig. 6C). The MLI analysis of the lungs stained with H & E indicate destruction in enlarged airspaces, indicating emphysematous injuries (Fig. 6 E & F).

**Figure 6.**
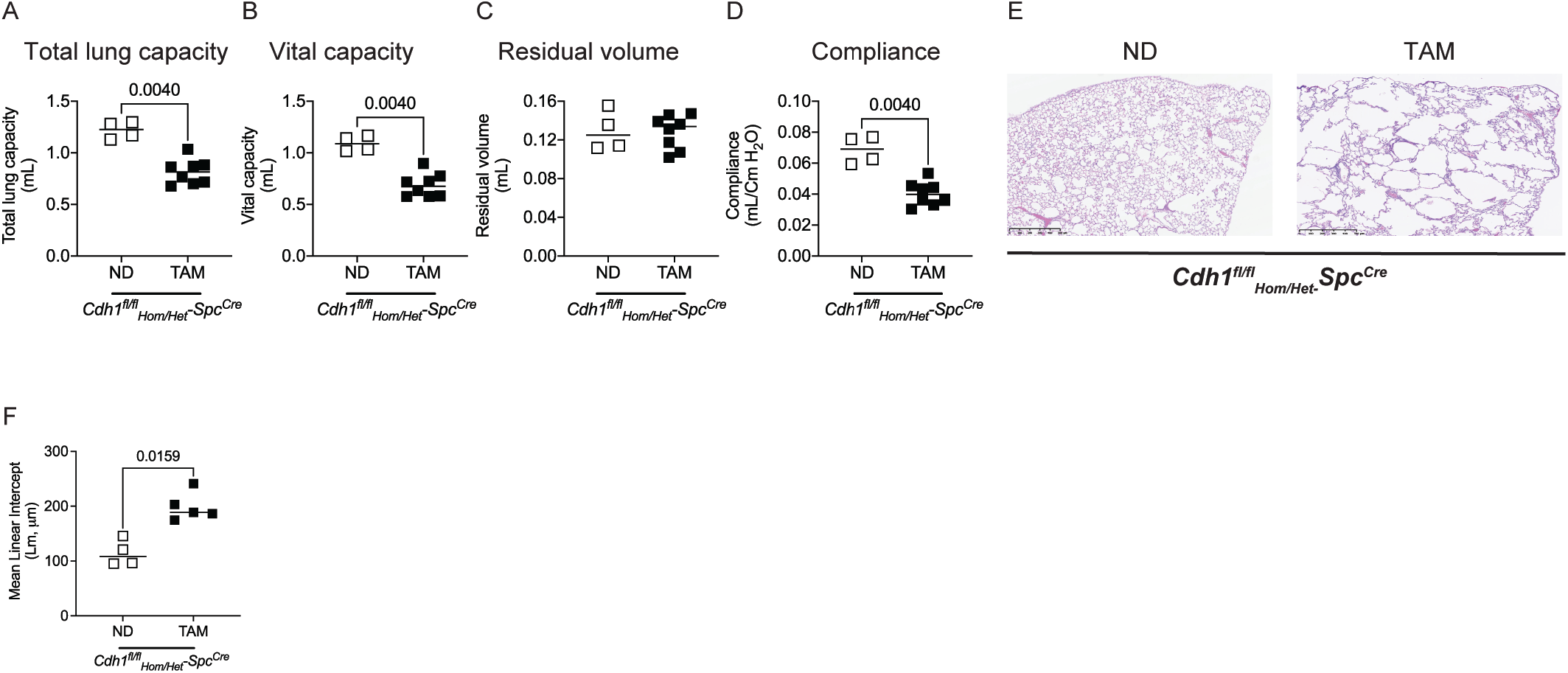
*In vivo* E-cadherin knockdown in the alveolar type II (AT2) cells of mice lung contributes to a increase in total lung capacity and compliance with a corresponding airspace enlargement. To knockdown E-cadherin in the AT2 cells of mice lung, *Cdh1*^*fl/fl*^_*Hom/Het*_*Spc*^*Cre*^ mice were fed tamoxifen (TAM) for 30 days. These were compared to *Cdh1*^*fl/fl*^_*Hom/Het*_*Spc*^*Cre*^ mice receiving normal chow diet (ND). Assessment of (A) total lung capacity, (B) vital capacity, (C) residual volume, and (D) compliance in AT2 cells of mice lung. (E) Representative H & E staining of mice lung parenchyma and (F) quantified mean linear intercepts (Lm) after E-cadherin knockdown in AT2 cells. Data is representative from 4 to 8 mice. *P* < 0.05 were considered statistically significant.

### Over expression of E-cadherin in COPD epithelia restores epithelial function

We further evaluated if restoring E-cadherin in COPD human bronchial epithelial (CHBEs) can restore the epithelial function. Increasing the E-cadherin expression in CHBEs with Ad5CMV-CDH1 significantly restored the barrier function and transited the epithelia from unjammed to jammed state (Fig. 7A – C). However, restoring E-cadherin, did not had any effect on % moving and CBF (Fig. 7D).

**Figure 7.**
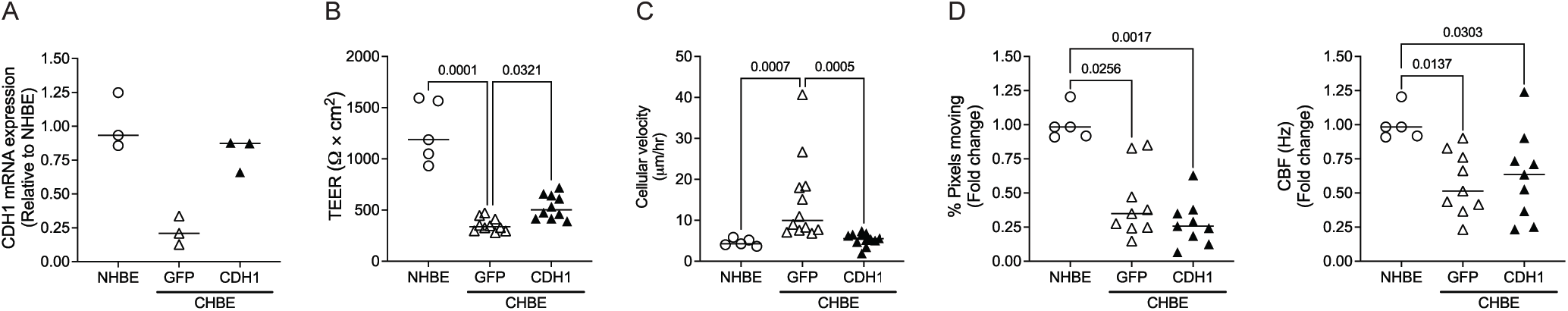
Over expression of E-cadherin in COPD epithelia (CHBE) restores epithelial function. CHBE at ALI were transfected Ad-GFP-U6-h-CDH1 (CDH1) to overexpress E-cadherin or Ad-GFP (GFP) as control at 2 × 10^9^ pfu mL^-1^. (A) Assessment of mRNA expression of *CDH1* in CHBE transfected with GFP/CDH1 compared to age and gender matched NHBE. Assessment of epithelial functional phenotypes such as (B) barrier function, (C) cellular velocity, and (D) percentage moving cilia (left panel) and ciliary beat frequency (right panel) of CHBE transfected with GFP/CDH1 were compared to non-diseased epithelium (NHBE). Data is representative of 2 donors and 2 to 3 inserts per donor. *P* < 0.05 were considered statistically significant. The data of NHBE is representation from Figure 8.

### Nrf2-activator has a protective role in epithelial plasticity and E-cadherin expression in COPD epithelia and cigarette-smoke (CS) exposed NHBE epithelia

To evaluate if a Nrf2 pathway activator as a specific pharmacological target, we treated the CHBE and CS exposed NHBE with CDDO-Me. The CDDO-Me significantly improved barrier function, pushes the epithelium to a jammed state and increased CDH1 mRNA expression (Fig. 8 A – C). Similarly, with CS injured epithelium, the CDDO-Me treatment significantly improved barrier function, allow for epithelial jamming and protected against CDH1 mRNA downregulation induced by CS (Fig. 8D – F). However, CDDO-Me had no protection against loss of ciliated cells as quantified by % pixels moving and CBF in both COPD and CS-injured epithelia (Fig. 8G & H).

**Figure 8.**
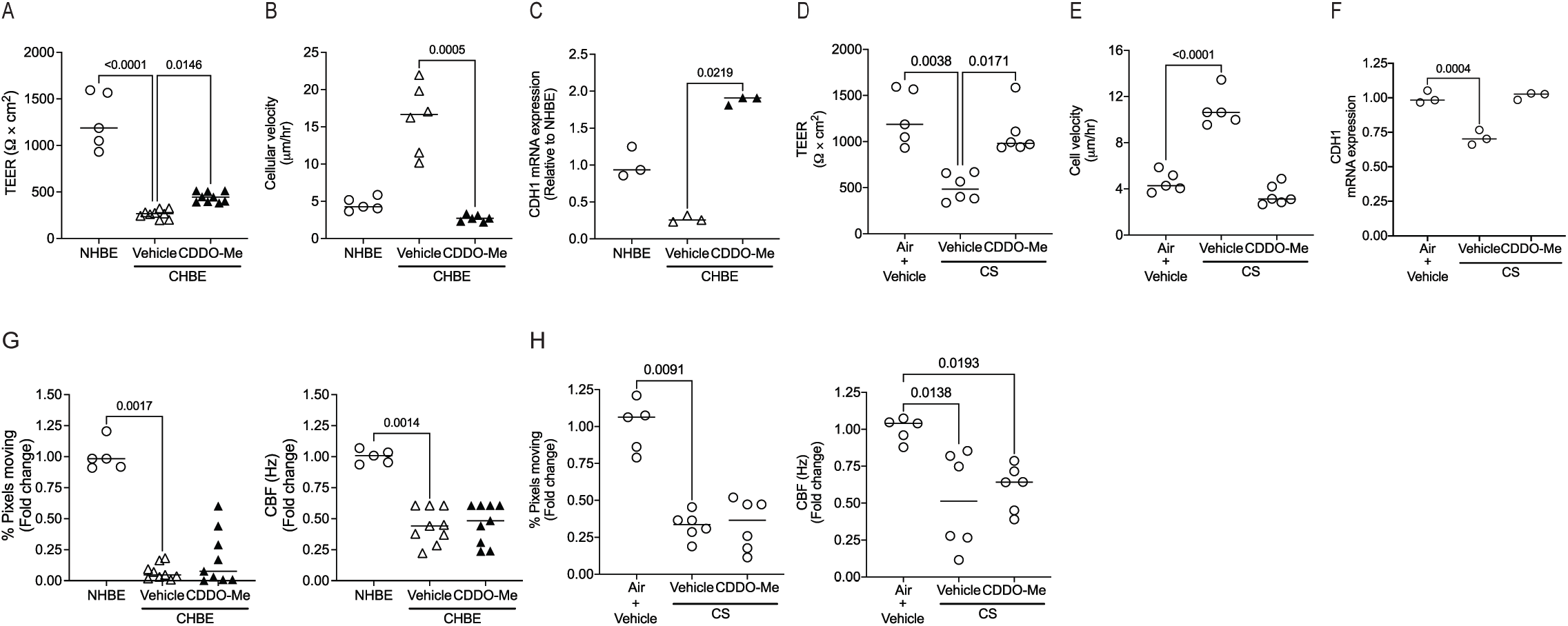
Nrf2 activator protects against E-cadherin expression and restores epithelial function in COPD epithelia (CHBE) and cigarette-smoke (CS) injured epithelia. The CHBE or the non-diseased epithelia (NHBE) exposed to CS were treated with 100 nM of CDDO-Me for 10 days. Assessment of CHBE (+/- CDDO-Me) epithelial functional phenotypes such as (A) barrier function, (B) cellular velocity and (C) CDH1 mRNA expression were compared to age matched non-diseased epithelium (NHBE). Assessment of CS exposed NHBE (+/- CDDO-Me) epithelial functional phenotypes such as (D) barrier function, (E) cellular velocity and (F) CDH1 mRNA expression were compared to NHBE exposed to air. Assessment of CHBE (+/- CDDO-Me) for (G) percentage moving cilia (left panel) and ciliary beat frequency (right panel) were compared to NHBE. Assessment of CS exposed NHBE (+/- CDDO-Me) epithelial functional phenotypes such as (H) percentage moving cilia (left panel) and ciliary beat frequency (right panel) were compared to NHBE exposed to air. Data is representative of 2 donors and 2 to 3 inserts per donor. *P* < 0.05 were considered statistically significant.

### Protective role of E-cadherin overexpression in cigarette-smoke injured epithelium

To evaluate if E-cadherin overexpression has a protective role against CS exposed epithelium. We observed that overexpression of E-cadherin in mTECs has a protective role against CS induced leakier epithelia (Fig. 9A), transition to a jammed epithelium (Fig. 9B), and protection against Cdh1 mRNA expression (Fig. 9C), without altering % pixels moving and CBF (Fig. 9D).

**Figure 9.**
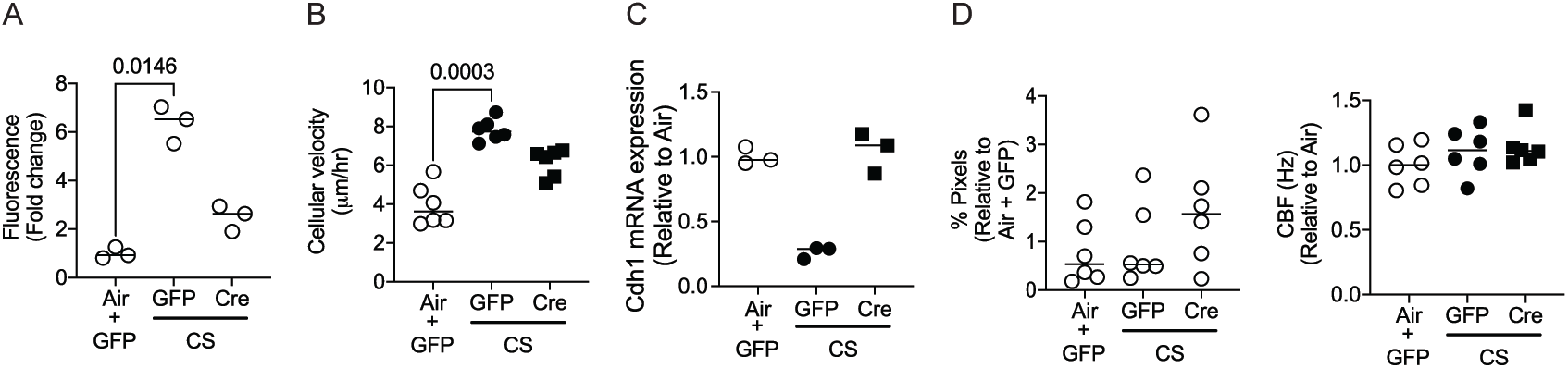
*In vitro* conditional E-cadherin knock-in has a protective role against cigarette-smoke (CS) injured mTECs. To induce the over-expression of E-cadherin in Cdh1 knock-in mice, mTECs were transfected with adeno-Cre (Ad5CMVCre-eGFP) instillation at 2 × 10^9^ pfu mL^-1^ and were exposed to CS for 10 days. Assessment of epithelial functional phenotypes such as (A) permeability, (B) cellular velocity, (C) *Cdh1* mRNA expression and (D) percentage moving cilia (left panel) and ciliary beat frequency (right panel), in Adeno-Cre (Cre) transfected epithelia were compared CS and air epithelia were transfected with Adeno-Ctrl (GFP). Data is representative of 6 mice and 3 to 6 inserts per mice. *P* < 0.05 were considered statistically significant.

## Discussion

E-cadherin is a principal component of the adherens junction and critical in maintaining cell-cell adhesion (18, 25–27). We and others have generated *in vitro*, animal, and human data indicating that loss of E-cadherin occurs after chronic tobacco exposure and in COPD patients (4–6, 16, 28). However, evidence that this loss of E-cadherin is not simply associated with disease but has a causal role in tissue remodeling in the adult lung and can serve as a therapeutic target to abrogate or reverse disease is lacking. The current study explores the *in vitro, ex vivo*, and *in vivo* models to study the effects of E-cadherin knockdown on lung epithelial function and architecture in both human and mouse models. While the entire lung epithelium is exposed to CS, we find an array of clinical manifestations in patients with COPD, the most common being emphysematous changes and evidence of chronic bronchitis. We observed that cell-specific loss of E-cadherin in the lung epithelium of the adult mouse results in various pathologic phenotypes seen in patients with COPD, with E-cadherin knockdown in ciliated cells causing increased airway hyperresponsiveness, and ATII epithelial cells causing emphysematous changes.

Recent data suggests that E-cadherin knock out from mice lungs *in utero* causes postnatal goblet hyperplasia and eosinophilia (23). While this was not found in loss of E-cadherin in the adult mouse, we found that loss of E-cadherin in ATII cells leads to increased lung volume and increased compliance, characteristics of emphysema which is in accords with Post *et al* (23). As a primary role of ATII cells are in cellular regeneration in the lung, these findings suggest that loss of E-cadherin impairs the regenerative capabilities of alveolar cells, which must be needed for maintenance of optimal lung volume and compliance.

While airway hyperresponsiveness is a characteristic feature in asthma, airway narrowing clearly occurs in patients with chronic bronchitis (29, 30). Mouse airways do not exhibit similar phenotypic changes in their airways that are seen in patients, so even showing increased reactivity to additional insults is indicative of changes in the airway with loss of E-cadherin in ciliated cells. We observed that knock down of E-cadherin in ciliated cells causes airway hyperresponsiveness without any changes in histology and pressure volume curve. These data coupled with the *in vitro* evidence of monolayer dysfunction with loss of E-cadherin seen in the NHBE cells suggest that specific loss of E-cadherin in the airway epithelium contributes to phenotypic changes that can occur with COPD.

Our findings of E-cadherin loss of AT1 cells are particularly intriguing. As ATI cells cover majority of alveolar surface(31–33), one could hypothesize that loss of E-cadherin would cause complete alveolar disruption with overwhelming injury. However, these mice were surprisingly minimally affected, raising the question of the relative contribution of E-cadherin to a mature alveolar monolayer. When AT1 epithelial cells are injured, AT2 epithelial cells undergo proliferation and differentiate into AT1 cells to regenerate the alveolar surface (34), and the data lead us to conclude that E-cadherin plays a more prominent role in this regenerating epithelium.

However, we do observe that knockdown of E-cadherin in AT1 epithelial cells increase mRNA expression of markers of epithelial to mesenchymal transition (EMT). EMT can result in fibrosis and these markers coupled with the increase in type III collagen, and the restrictive lung defects identified in the mouse model suggests fibrosis. We have previously reported that with repetitive injury to airway epithelium with cigarette smoke the loss of E-cadherin is coupled with an increase in markers of EMT (4). Most studies suggested that loss of E-cadherin and concomitant upregulation of N-cadherin (neural cadherin) expression is the hallmark of EMT, though whether loss of E-cadherin is necessary and sufficient to induce EMT has been of debate (35, 36). While alveolar fibrosis is not associated with COPD, there are other tobacco smoking has been associated with other fibrotic lung diseases such as Idiopathic Pulmonary Fibrosis and even Respiratory bronchiolitis-interstitial lung disease (RB-ILD). While E-cadherin loss and EMT have been implicated in some studies (37), the data are mixed and less conclusive (27) and further studies are needed to tease the relative contribution of E-cadherin loss to these diseases.

In conclusion, for the first time we have evidence that loss of E-cadherin is indeed causal to the pathology seen in patients COPD, with cell-specific losses contributing to different disease phenotypes. While there are currently few strategies that are designed to target E-cadherin and monolayer integrity for therapeutic potential, our study provides evidence that such a strategy would indeed have merit.

## Methods

### Human airway epithelial cell culture

Primary non-diseased human bronchial epithelial (NHBE) and COPD human bronchial epithelial (CHBE) cells were purchased from MatTek Life Sciences (Ashland, MA, USA) or Lonza (Basel, Switzerland) were expanded on collagen-coated T_75_ flask and differentiated into pseudostratified epithelium at air-liquid interface (ALI) on 0.4 µm pore size polyester membrane Transwells as described by us before (4–6).

### E-cadherin knockdown in NHBEs

Adenovirus to knock out E-cadherin (Ad-GFP-U6-h-CDH1-shRNA) and Control Adenovirus-based scrambled shRNA with GFP (Ad-GFP-U6-shRNA, GFP), were purchased from Vector Biolabs (Malvern, PA, USA) to transduce cells. Well-differentiated NHBE at 4 to 6 weeks ALI was treated apically and basolateral with transfection media (containing PneumaCult™-ALI medium and 5 μg mL^-1^polybrene with Ad-GFP-U6-h-CDH1-shRNA or Ad-GFP at 3 × 10^9^ pfu mL^-1^) for 4 hours. After 4 hours, the transfection media was transferred from apical surface to basolateral. After 20 hours, the transfecting media was completely removed and replaced with PneumaCult™-ALI medium media. The culture was maintained for additional 48 hours. Post 48 hours, the epithelium at ALI was visualized for Green Fluorescent Protein (GFP) at 20X by fluorescent microscopy using 3i Marianas/Yokogawa Spinning Disk Confocal Microscope (Leica Microsystems, TX, USA), assessed for functional phenotypes (as described below), and samples were collected for western blot assay.

### E-cadherin overexpression in CHBEs

Adenovirus to overexpress E-cadherin (Ad-GFP-U6-h-CDH1) and control adenoviruses (Ad-GFP), were purchased from Vector Biolabs (Malvern, PA, USA) to transduce CHBE at 4 to 6 weeks ALI at 2 × 10^9^ pfu mL^-1^ as described in earlier sections of E-cadherin knockdown in NHBEs.

### Cigarette-smoke (CS) exposure to NHBEs

The NHBEs at 4 to 6 weeks ALI were exposed to either exposed to CS smoke or humidified air for 10 days as mentioned previously by us(4–6). One CS exposure consisted of 2 cigarettes which burned for ∼ 8 minutes using the ISO puff regimen.

### Treatment of Nrf2-pathway activator

In this study CDDO-Me (Toronto Research Chemicals) a potent Nrf2-pathway activator at a dose of 100 nM was utilized. The basolateral treatment with CDDO-Me was 5 days for CHBE and 10 days for the NHBE at ALI exposed to CS.

### Quantification of epithelial functional phenotypes

The epithelial functional phenotypes were assessed by quantifying the barrier function, paracellular permeability, ciliary beat frequency (CBF), % moving cilia and cellular velocity as described below.

### In vitro barrier function assessment

To determine epithelial integrity of the human and mice pseudostratified monolayer epithelium at ALI, the trans-epithelial electrical resistance (TEER) was measured using epithelial voltohmeter (EVOM, World Precision Instruments Inc, FL, USA) with the STX2 electrodes as previously described (4–6). Values were corrected for fluid resistance (insert with no cells) and surface area.

### Paracellular permeability

The paracellular permeability of the epithelium at ALI was determined using fluorescein isothiocyanate-dextran (FITC-Dextran) flux assay as described previously (4, 5).

### CBF

The epithelium at ALI were incubated at 37°C and 5% CO_2_ in the 3i Marianis/Yokogawa Spinning Disk Confocal Microscope (Leica Microsystems, TX, USA) and high speed time-lapse videos were taken at 32X air at 100 Hz with a total of 250 frames using a scientific Hamamatsu C1140-42U30 CMOS camera (Hamamatsu Photonics, NJ, USA) as reported in our previous publications (4, 6). Five areas were imaged per insert. A Matlab (R2020a) script (validated against SAVA (38)) was used to determine average CBF and % moving cilia per video to generate a heatmap(39).

### Cellular velocity

Cellular velocity was quantified by performing Particle Image Velocimetry (PIVlab) on Matlab (R2020a), using a multi-pass cross correlation analysis with decreasing interrogation window size on image pairs to obtain the spatial velocity as described previously by us (40). Using a phase contrast microscopy at 3i Marianis/Yokogawa Spinning Disk Confocal Microscope at 32X, and 37°C and 5% CO_2_ incubation, we captured time-lapse videos of the pseudostratified monolayer epithelium for every 5 minutes for 2 hours and average velocity was computed for the area. Five areas were imaged per insert.

### Animal and study design

The study was approved by the Institutional Animal Care and Use Committee (IACUC) of the Johns Hopkins University Animal Use and Care Committee and compiled with the Guidelines for Care and use of Laboratory Animals issued by the USA National Institute of Health. The study consisted of *Cdh1*^*fl/fl*^, *Foxj1-CreER*^*T2*^, *Sftpc-CreER*^*T2*^, and *Ager-CreER*^*T2*^. The *Cdh1*^*fl/fl*^, *Foxj1-CreER*^*T2*^, and *Ager-CreER*^*T2*^ were purchased from The Jackson Laboratory (Bar Harbor, ME). The mating pair for *Sftpc-CreER*^*T2*^ mice (MGI ID: 5444645) were provided by Dr. Harold A Chapman (Lung Biology Center, UCSF, San Francisco) (41). All mice were bred and maintained in a specific pathogen-free environment.

To generate E-cadherin specific knock out in ciliated cells, *Cdh1*^*fl/fl*^ were crossed with *Foxj1-CreER*^*T2*^ mice, to generate tamoxifen responsive *Cdh1*^*fl/fl*^(*Foxj1-CreERT*^*2*^) litters. To generate E-cadherin specific knock out in alveolar type I cells, *Cdh1*^*fl/fl*^ were crossed with *Ager-CreER*^*T2*^ mice, to generate tamoxifen responsive *Cdh1*^*fl/fl*^(*Ager-CreER*^*T2*^) litters. To generate E-cadherin specific knock out in alveolar type II cells, *Cdh1*^*fl/fl*^ were crossed with *Sftpc-CreER*^*T2*^ mice, to generate tamoxifen responsive *Cdh1*^*fl/fl*^ (*Sftpc-CreERT*^*2*^) litters. The mice were genotyped according to the optimized PCR primers as described in Table S1.

We have created a mouse Cdh1 conditional knock-in (gene for E-cadherin) at the locus of ROSA26 in C57BL/6 mice by CRISPR/Cas-mediated genome engineering. The CAG-loxP-Stop-loxP-Kozak-mouse Cdh1 CDS-polyA cassette was cloned into intron 1 of ROSA26 in reverse orientation. The expression of mouse *Cdh1* is dependent on Cre recombinase.

### Tamoxifen administration

Beginning at 5 weeks of age, mice were switched to either tamoxifen chow diet (TD.130856, Envigo, IN, USA) or normal chow diet (Envigo, IN, USA) for a period of 31 days.

### Intratracheal instillation to knock down E-cadheirn in mice lung

*Cdh1*^*fl/fl*^ mice were anesthetized with ketamine (100 mg/kg) - xylazine (15 mg/kg) solution. To knock out E-cadherin from *Cdh1*^*fl/fl*^ mice, lung instillation of either Adenovirus Ad-5CMVeGFP (Control) and Ad5CMVCre-eGFP (Cre) at titer of 2×10^9^ pfu in 60 μL was performed as described previously on 10^th^ day for a period of 1 month or 3 months (42).

### Physiological measurements

Mice were anesthetized with ketamine (100 mg/kg) - xylazine (15 mg/kg) solution. Once sedated, the trachea was cannulated with 18-gauge needle. Electronic controls on the ventilator are set to periodically lengthen the inspiratory time to provide a measurement of total lung capacity, vital capacity, respiratory resistance, and lung compliance. The *Cdh1*^*fl/fl*^(*Foxj1-CreERT*^*2*^) mice were further given stepwise increases (0, 3 & 30 mg/mL) of Acetyl- β- methylcholine chloride (Sigma Aldrich, MO, USA), to assess airway responsiveness.

### Lung fixation and histopathology

After performing physiological measurements, the lungs were fixed by inflation with 25cmH_2_0 with instilled formaldehyde and fixed lung volume were assessed as described previously by us (42). The fixed lungs were embedded in paraffin and sagittal sections were stained with hematoxylin and eosin (H&E) at Reference Histology Laboratory, Johns Hopkins Medical Institutions – Pathology (Baltimore, USA).

### Isolation of mice tracheal epithelial cells (mTECs)

The *Cdh1*^*fl/fl*^ mice were euthanized by following Carbon dioxide narcosis followed by cervical dislocation IACUC guidelines. The tissue free lumen exposed trachea dissected from the 10 *Cdh1*^*fl/fl*^ mice were added to 10 mL of 1X-Phosphate Buffer Saline (1X-PBS, ThermoFisher Scientific, New York, USA) supplemented with Penicillin-Streptomycin (ThermoFisher Scientific, NY, USA). The trachea was transferred to 15 mL 0.15% Pronase solution and incubated overnight at 4°C. The tube was rocked for 40-50 times and were passed through a 70 µm cell strainer (Corning Life Sciences, MA, USA). The cell strainer consists of digested trachea and the solution consists of cells. The cell solution was topped up to a volume of 50 mL with1X-PBS and centrifuged at 300 g with brake on. The resulting pellet was suspended with 15 mL of DMEM supplemented with 10% Fetal Bovine Serum (FBS) and Penicillin-Streptomycin and incubated at 37°C and 5% CO_2_ for 4 hours in a T_75_ flask (Corning Life Sciences, MA, USA) to eliminate fibroblasts and mononuclear cells (as fibroblasts and mononuclear cells get adhere to flask). After 4 hours, the cells were suspended in epithelial expansion media (Prepared by mixing 490 mL PneumaCult™-Ex Plus Basal Medium Supplemented with 10 mL PneumaCult™-Ex Plus 50X Supplement, 0.5 mL Hydrocortisone stock solution and 5 mL of 1% Penicillin-Streptomycin: StemCell Technologies Inc., Vancouver, Canada). The suspended cells are expanded in a T_75_ collagen-coated flask at 80-90% confluency. The cells were plated onto Rat tail collagen I coated 0.4 µm pore polyester membrane 12 mm Transwell® at a seeding density of 300,000 cells/well with epithelial expansion media on apical and basolateral surface. At 100% confluency, the transwells were put at air-liquid interface (ALI) media (Prepared by 450 mL PneumaCult™-ALI medium supplemented with 50 mL of PneumaCult™-ALI 10X Supplement, 5 ×1 mL vial of 100X PneumaCult™-ALI Maintenance Supplement, 1 mL Hydrocortisone Solution, 1 mL Heparin Solution and 5 mL of 1% Penicillin-Streptomycin; StemCell Technologies Inc., Vancouver, Canada) on the basolateral surface for 2 weeks to obtain a fully differentiated epithelium.

### In vitro knock down of E-cadherin in mTECs

The mTECs of *Cdh1*^*fl/fl*^ mice at 2 weeks ALI were transfected with either Adenovirus Ad-5CMVeGFP (Control) or Ad5CMVCre-eGFP (Cre) at titer of 2×10^9^ pfu mL^-1^ with transfection protocol as described in earlier sections of E-cadherin knockdown in NHBEs.

### Over-expression of E-cadherin in mTECs

The mTECs of *Cdh1* conditional knock-in mice at 2 weeks ALI were transfected with either Adenovirus Ad-5CMVeGFP (Control) or Ad5CMVCre-eGFP (Cre) at titer of 2×10^9^ pfu mL^-1^ with transfection protocol as described in earlier sections of E-cadherin knockdown in NHBEs.

### Western blot assay

Equal quantities of protein were separated on NuPAGE™ 4 – 12%, Bis-Tris gradient gel (ThermoFisher Scientific, NY, USA) using a Mini Gel Tank (ThermoFisher Scientific, New York, USA) at 200V for 30 minutes. Proteins were transferred to a Immobilion-P PVDF membrane (Millipore Sigma, MA, USA) at 47V overnight at 4°C using Mini Trans-Blot® Cell (Bio-Rad Laboratories, CA, USA). Following the transfer, the blots were blocked with 5% w/v of Non-Fat Powdered Milk (Boston BioProducts, MA, USA) in 1X PBS with 0.1% Tween® 20 Detergent, (1X PBST, Millipore Sigma, MA, USA) for 2h at room temperature on a shaker. The blocked membrane was washed with 1X PBST and then incubated in primary antibody (E-cadherin (24E10) Rabbit mAb, 135 kDa and GAPDH (14C10) Rabbit mAb, 37 kDa antibodies from Cell Signaling Technology, MA, USA) at a concentration of 1:1,000 overnight at 4°C. The blot was further washed with 1X PBST and then incubated in secondary antibody IRDye® 800CW Goat anti-Rabbit (LI-COR, NE, USA) at a dilution of 1:10,000 for 1h at room temperature. Blots were washed 3 times with 1X PBST, then visualized on a Li-Cor imager. Images were analyzed using Fiji (43).

### Immunofluorescence assay

The lung tissue sections were prepared for immunofluorescent staining as described by Bratthauer (37). The tissue sections were permeabilized with 0.1% Triton™ X-100 (Millipore Sigma, MO, USA) in 1X PBS and blocked for 2 hours in 1X PBS with Normal goat serum (10%) (ThermoFisher Scientific, MA, USA). The primary antibody (E-cadherin (24E10) Rabbit mAb) diluted with 1:200 in blocking solution, was incubated on the tissue slides for overnight at 4°C. The tissue sections were further washed with 1X PBS and incubated with Goat anti-Rabbit IgG (H+L), Super clonal Recombinant Secondary Antibody, Alexa Fluor 555 (ThermoFisher Scientific, MA, USA) diluted at 1:10,000 for 1 hour at room temperature. The tissue was then stained with 2 µg mL^-1^ of Hoechst 33342 (ThermoFisher Scientific, MA, USA) for 15 minutes at room temperature, washed, and mounted. The images were then captured in Zeiss LSM700 Confocal microscope.

### Quantitative polymerase chain reaction (qPCR) assay

Total RNA was isolated from cultured from human bronchial epithelial cells / mice right lung tissues and purified using the Qiagen AllPrep DNA/RNA Mini Kit (Qiagen, Hilden, Germany), supplemented with the Proteinase K (Qiagen, Hilden, Germany). cDNA of 1000 ng µL^-1^ was obtained using the High-Capacity cDNA Reverse Transcription Kit (ThermoFisher Scientific, MA, USA), and the absence of DNA contamination was verified by excluding the reverse transcriptase from subsequent PCR reaction. cDNA was subjected to PCR using the SYBR™ Green PCR master Mix (ThermoFisher Scientific, MA, USA) to amplify the epithelial, or mesenchymal or fibrosis markers listed in supplementary Table S2.

Each PCR reaction was carried out as follows: initial denaturation of 94°C for 15 minutes, 45 cycles of 94°C for 35 seconds, 60°C for 1 minute and 72°C for 1 minute 15 seconds, followed by a final extension at 72°C for 2 minutes. Each cycle was repeated 45 times. Based on comparative Ct method, gene expression levels were calculated and GAPDH was used as housekeeping gene.

### Statistics

Prism version 9.0 (GraphPad, CA, USA) and Matlab (R2020a) was used to analyze the data. The results are expressed as a median. To compare the data sets involving 2 groups, Mann-Whitney test was performed. To compare more than 2 groups, Kruskal-Wallis test followed by Dunn’s multiple comparison test was performed.

## Acknowledgements

We would like to thank Johns Hopkins University (JHU) – School of Medicine (SoM) Microscope Facility for providing access and training in 3i Marianas Spinning Disk Confocal and Zeiss LSM700 single-point laser scanning confocal microscope. We would also like to thank JHU-SoM Reference Histology Laboratory for assisting in H & E and Masson trichrome staining of mice lung tissues. The Research reported in this publication was supported by the National Heart, Lung, and Blood Institute (NHLBI R01-HL124099 and HLR01-HL151107), and Office of the Director of the National Institutes of Health under award number S10 OD016374 (to Scot Kuo of the JHU Microscope Facility).

