## Supplementary material for "Knockout of E-cadherin in adult mouse epithelium results in emphysema and airway disease"

**Supplementary Information**

*Table S1: Primer details for genotyping the Mice*

| **Gene** | **Primer (5’-3’)** | **Product details** |
| --- | --- | --- |
| *Cdh1*  *(E-cadherin)* | P1: GGG TCT CAC CGT AGT CCT CA  P2: GAT CTT TGG GAG AGC AGT CG | Mutant – 310 bp  Heterozygote – 243 bp & 310 bp  Wild type – 243 bp |
| *Foxj1*  *(Ciliated cells)* | P1: GCA GAT GGA GAG AGG TGG AG  P2: CTT GGC GTT GAG AAT GGA GA  P3: ATT GCA TCG CAT TGT CTG AG | Mutant – 294 bp (P2 & P3)  Heterozygote – 294 bp & 472 bp  Wild type – 472 bp (P1 & P2) |
| *Ager*  *(Alveolar Type I cells)* | P1: ATC GCA TTC CTT GCA AAA GT  P2: GGA CTC TTG TCC CAG AAG CA | Mutant - ~500 bp  Heterozygote - ~500bp & 204 bp  Wild type – 200 bp |
| *Spc*  *(Alveolar Type II cells)* | P1: ATG TCC AAT TTA CTG ACC G  P2: CGC GCC TGA AGA TAT AGA AG | ~250 bp |

*Table S2: qPCR primer details to evaluate EMT and fibrosis markers*

| **Gene** | **Primer** | **Product details** |
| --- | --- | --- |
| ***Human*** |  |  |
| *CDH1* | F: GCC TCC TGA AAA GAG AGT GGA AG  R: TGG CAG TGT CTC TCC AAA TCC G | 131 bp |
| *GAPDH* | F: GTC TCC TCT GAC TTC AAC AGC G  R: ACC ACC CTG TTG CTG TAG CCA A | 131 bp |
| ***Mouse*** |  |  |
| *Cdh1* | F: CAT CAC TGC CAC CCA GAA GAC TG  R: ATG CCA GTG AGC TTC CCG TTC AG | 153 bp |
| *Cdh2* | F: CCT CCA GAG TTT ACT GCC ATG AC  R: GTA GGA TCT CCG CCA CTG ATT C | 149 bp |
| *Col3a1* | F: GAC CAA AAG GTG ATG CTG GAC AG  R: CAA GAC CTC GTG CTC CAG TTA G | 114 bp |
| *Gapdh* | F: CAT CAC TGC CAC CCA GAA GAC TG  R: ATG CCA GTG AGC TTC CCG TTC AG | 153 bp |
| *Slug/Snai2* | F: TCT GTG GCA AGG CTT TCT CCA G  R: TGC AGA TGT GCC CTC AGG TTT G | 133 bp |
| *Snai1* | F: TGT CTG CAC GAC CTG TGG AAA G  R: CTT CAC ATC CGA GTG GGT TTG G | 163 bp |
| *Twist1* | F: GAT TCA GAC CCT CAA ACT GGC G  R: AGA CGG AGA AGG CGT AGC TGA G | 134 bp |
